# Transcriptome analysis of the tardigrade *Hypsibius exemplaris* exposed to the DNA-damaging agent bleomycin

**DOI:** 10.1101/2024.02.01.578372

**Authors:** Yuki Yoshida, Akiyoshi Hirayama, Kazuharu Arakawa

**Affiliations:** Institute for Advanced Biosciences, Keio University, Tsuruoka, Yamagata, 997-0017, Japan; Graduate School of Media and Governance, Keio University, Fujisawa, Kanagawa, 252-0882, Japan; Faculty of Environment and Information Studies, Keio University, Fujisawa, Kanagawa, 252-0882, Japan; Exploratory Research Center on Life and Living Systems (ExCELLS), National Institutes of Natural Sciences, Okazaki, Aichi, 444-8787, Japan

**Keywords:** Tardigrade, Bleomycin, Transcriptome, Metabolome, Tryptophan

## Abstract

Tardigrades are microscopic animals that are renowned for their capabilities of tolerating near-complete desiccation by entering an ametabolic state called anhydrobiosis. However, many species also show high tolerance against radiation in the active state as well, suggesting cross-tolerance via the anhydrobiosis mechanism. Previous studies utilized indirect DNA damaging agents to identify core components of the cross-tolerance machinery; however, it was difficult to distinguish whether transcriptomic changes were the result of DNA damage or residual oxidative stress. To this end, we performed transcriptome analysis on bleomycin-exposed *Hypsibius exemplaris*. We observed induction of several tardigrade-specific gene families that may be the core components of the cross-tolerance mechanism. We also identified an enrichment of the tryptophan metabolism pathway, which metabolomic analysis suggested the engagement of this pathway in stress tolerance. These results provide several candidates for the core component of the cross-tolerance, as well as possible anhydrobiosis machinery.

## 1. Introduction

Tardigrades are microorganisms capable of entering an ametabolic state known as anhydrobiosis when faced with environmental desiccation ^1)^. In this state, tardigrades are able to tolerate extreme environments, ranging from low to high temperatures ^2–4)^, pressure ^5)^, organic solvents ^6)^, and high dosage of ultraviolet ^7, 8)^ or radiation ^9–17)^. Current knowledge of the anhydrobiosis machinery is consists of both mechanisms that are broadly conserved across eukaryotes and tardigrade-specific proteins ^18)^. Protection from potential damage during desiccation has been the focus of anhydrobiosis studies, and anhydrobiosis is known to sometimes cause DNA double strand breaks in tardigrades ^19)^ and also in sleeping chironomids ^20)^. One example is the nucleus-localizing protein Damage suppressor (Dsup), which has been identified in *R. varieornatus* and *H. exemplaris* to have DNA protective capabilities ^21–25)^, as expressing human cell lines have increased X-ray tolerance. However, the persistence of DNA damage in recovered samples suggests that the protection alone is not sufficient.

Interestingly, studies have observed high tolerance to gamma radiation in the active state in many species throughout the phylum Tardigrada, sometimes even higher than during anhydrobiosis, suggesting that these organisms have high DNA repair capabilities, possibly a cross-tolerance acquired through anhydrobiosis mechanisms ^26)^. The core machinery of tardigrade DNA repair does not differ significantly from that of other eukaryotes ^27)^, with the exception of the duplication of several genes in *Ramazzottius varieornatus*, e.g. *MRE11* ^28)^. Other DNA repair factors regulated during anhydrobiosis include the regulation of RAD51 in *Milnesium* cf. *tardigradum* ^29)^. Transcriptome-based studies have identified both non-homologous end joining repair and homologous recombination repair as regulated during anhydrobiosis ^30, 31)^. Other studies have also observed regulation of DNA damage repair pathways in specimens exposed to ionizing and non-ionizing radiation; gamma rays and ultraviolet C ^8, 32, 33)^. In addition, two novel genes families related to stress response have been identified: Anhydrobiosis related Mn (manganese) dependent Peroxidase (AMNP), a novel Mn^2+^ dependent peroxidase widely conserved throughout the phylum Tardigrada, and the Rv.g241 gene family, a potentially horizontally transferred gene with similarity to the C-terminus of bacterial peroxidases. However, the mechanism of anhydrobiosis is a complex union of various pathways, and the DNA damage caused under the conditions used above is mostly based on the indirect attack of reactive oxygen species, making it difficult to classify whether the differentially expressed genes were regulated as a response to DNA damage or oxidative stress. Therefore, a more focused approach was required to identify key factors in the tardigrade DNA repair process.

To this end, we performed a transcriptomic analysis of *Hypsibius exemplaris* exposed to bleomycin, a commonly used compound that induces DNA double strand breaks. *H. exemplaris* is a relatively weak anhydrobiote that requires 48 h of preconditioning at high relative humidity for successful anhydrobiosis ^34)^. We have previously obtained the genome of this species to identify key factors of anhydrobiosis ^31)^, and genetic toolkits (*e.g.*, RNAi) have also been developed for this species ^35)^. *H. exemplaris* shows tolerance to gamma rays in both embryonic and adult active stages at extremely high doses (3,000-4,000 Gy of 137Cs gamma ray) ^12)^, suggesting the existence of a prominent DNA repair protection, response, or repair machinery. The drug bleomycin is known to cause both direct and indirect DNA damage and would therefore be comparable to other DNA damaging stresses^36)^. We observed a three-tiered response, of which the early response had the most commonalities with anhydrobiosis entry through extensive time-course transcriptome sequencing. We found that several early responders that were observed in our previous analysis were included in the bleomycin early response, highlighting the importance of these gene families. We observed enrichment of the tryptophan metabolism pathway in the early response, but the metabolomic analysis was not consistent with the transcriptomic response. Taken together, we propose that the gene set identified here may comprise the core gene set required for cross-tolerance in *H. exemplaris*.

## 2. Results

### 2.1. The transcriptional response against bleomycin exposure

We generated 6-11M RNA-Seq reads (average 8.7M reads) for each sample, including an unexposed control sample, all in triplicates. Approximately 66-85% (average 79%) mapped to coding sequences (**Supplementary Table S1**). Approximately 70% of the transcripts had an average TPM greater than 1, consistent with previous studies. Initial clustering based on PCA analysis indicated that the transcriptome changed dynamically between 0-1h and 12-24h after bleomycin treatment (**Supplementary Figure 1AB**). We detected 11,210 transcripts that were differentially expressed between any time points. This is approximately three times more than our previous observations in anhydrobiosis, suggesting a dynamic shift in the cellular state during recovery from the 24h bleomycin exposure. To obtain an overview of the transcriptional response, we clustered the transcriptome profiles of differentially expressed genes (DEGs), resulting in eight groups (**Supplementary Figure 1B**). Gene Ontology enrichment analysis of each group (**Supplementary Table S2**) showed that terms related to DNA repair were enriched in Groups 3, 5, 8, of which only Group 5 contained terms related to double strand break repair, in particular non-homologous end joining. Groups 3 and 8 were enriched for mismatch repair and nucleotide-excision repair.

These eight clusters could be grouped into three larger clusters, designated Clusters Early (E; Groups 2, 3, 8), Middle (M; Groups 4, 5), and Late (L; Groups 1, 6, 7) (**Supplementary Figure 1C**). Gene ontology enrichment analysis of these groups suggested a three-step regulation of cellular processes (**Figure 1**). First, stress-response-related metabolism, DNA synthesis, and cellular signaling occur in the early response (1-3 h), possibly preparing to counteract the oxidative stress and DNA damage produced by bleomycin. These processes are followed by mitochondrial regulation, indicating recovery from mitochondrial stress (3-12 hours). Finally, DNA replication and metabolism are regulated in the late stages to restore homeostasis. We also validated the expression of previously identified tardigrade-specific anhydrobiosis-related genes, such as *CAHS*, *SAHS*, *MAHS*, *LEAM*, *Dsup*, *TDR1*, *AMNP*, and *Rv.g241* gene families ^22, 33, 37–39)^. In contrast to previous studies using stresses other than anhydrobiosis, most CAHS and SAHS copies showed a significant increase in expression. On the contrary, the expression of MAHS (BV898_03788) and Dsup (BV898_01301) orthologs was significantly decreased during the time course. The LEAM ortholog (BV898_03429) did not show any significant change. The tardigrade-specific peroxide AMNP family and the stress-responsive Rv.g241 gene family also showed an increase in expression (**Supplementary Figure S1DE**). We observed extremely high expression of TDR1 at 0h (TPM 328), of which approximately TPM 100 is observed until 12h.

**Fig. 1.**
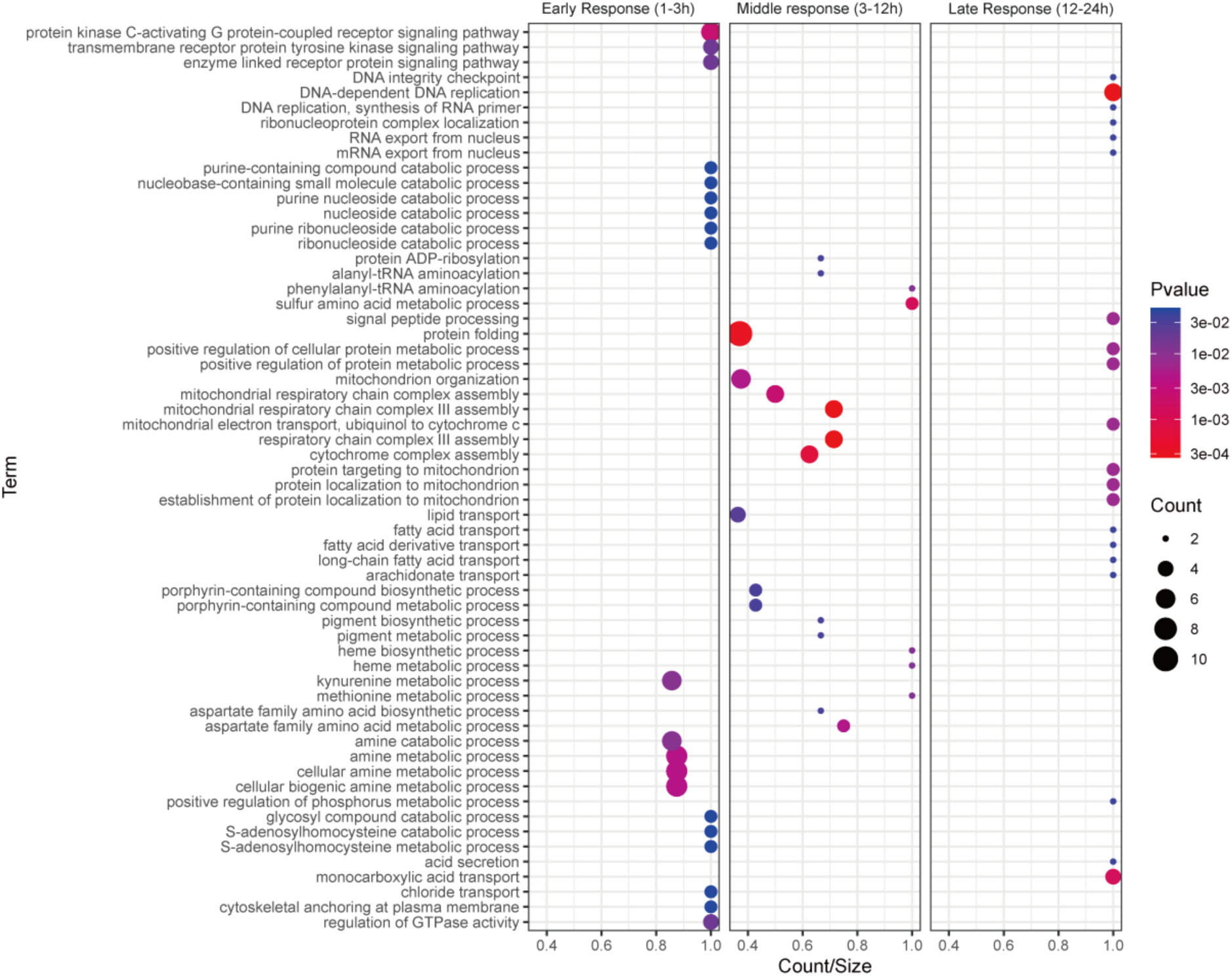
Gene ontology enrichment analysis of Early, Middle, and Late responses. Transcripts with top 20 odd ratios are shown. The X-axis indicates the number of transcripts with the corresponding terms in DEGs (Count) divided by the number in the whole genome (Size). Y-axis indicates the Biological Process. Dot colors and sizes indicate the p-value and count, respectively.

### 2.2. Comparison with anhydrobiosis entry and identification of novel stress-responsive transcripts

We then compared the bleomycin response with anhydrobiosis entry. Re-analysis of the anhydrobiosis transcriptome data revealed 2,738 DEGs, of which 2,089 transcripts were significantly upregulated. The anhydrobiosis entry shared the largest number of transcripts with the Early response cluster (1,152), followed by the Late cluster (692) and then the Middle cluster (449) **(Figure 2A**), suggesting that the Early response may partially resemble the anhydrobiosis entry trajectory. To obtain an overview of the transcripts common to anhydrobiosis entry and each response group, we subjected each subgroup to Gene Ontology enrichment analysis (**Supplementary Table S3, Supplementary Figure S2ABC**). We found that most of the enriched terms were time specific (37, 16, and 31 terms for Early, Middle, and Late response, respectively, **Figure 2B**), but the terms “proteolysis” and “oxidation-reduction process” were common to all subgroups, emphasizing the importance of antioxidative stress processes. We then focused on the 37 terms common to the Early response and anhydrobiosis, and interestingly found many terms related to transmembrane transport and the tryptophan metabolism to be contained in this list.

**Fig. 2.**
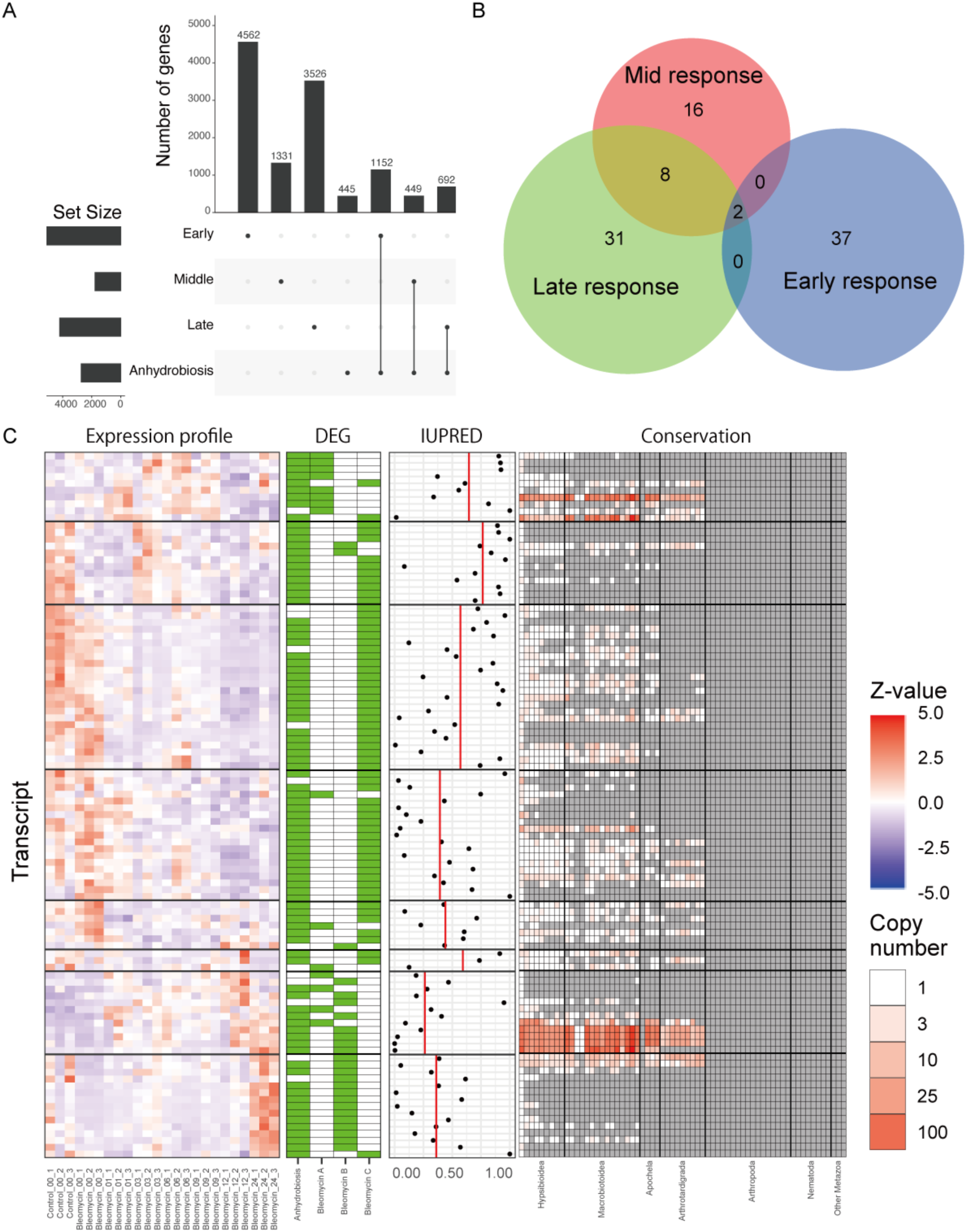
Comparison with anhydrobiosis entry. [A] UpSet plot showing the number of genes shared between each cluster and anhydrobiosis entry. [B] Number of GO BP terms significantly enriched between each response cluster and anhydrobiosis. [C] Features of the 102 NLS predicted transcripts without annotations, induced in anhydrobiosis and either one of lusters A, B, C. Transcripts are ordered based on the clustering preformed Figure S1C. Z-scaled expression profiles, DEG profiles, IUPRED3 score, conservation patterns across Tardigrada and relative lineages are shown.

We next screened the DEGs to attempt to identify novel stress-responsive transcripts without SwissProt orthologs. First, we aimed to obtain Early responding gene families, induced in anhydrobiosis entry and bleomycin early-response that have orthologs induced in *R. varieornatus* UVC exposure early-response (**Supplementary Table S4**). Under these thresholds, we identified 9 unannotated gene families, including two CAHS families (OG0001789, OG0002138), the AMNP and Rv.g241 gene families (OG0000230, OG0000231). For the remaining gene families, protein structures were predicted with AlphaFold2 using ColabFold and similarity search with DALI against the AlphaFold database (**Supplementary Table S4**). We found protein families with similarity (Identity > 30 or Z-score > 10) to metalloendopeptidases (OG0000456), hormone receptors (OG0000592), abnormal cell migration proteins (OG0000940), and LYSM domain proteins (OG0028858).

Only one protein family could not be annotated (OG0010176). Additional screening with fold change (Max TPM / Control) > 2 and high expression (Max TPM>100) left only the CAHS, Rv.g12777, and Rv.g241 gene families, again, highlighting the importance of these gene families. The TDR1 gene did not have a significant increase in expression in our anhydrobiosis dataset (TPM 0.9 to 6.19, 6.87x increase, FDR=0.36).

We also tested for nucleus-localizing stress-responsive proteins: tardigrade specific proteins with predicted nuclear localization signals (NLSs) and induced by both anhydrobiosis and bleomycin (**Figure 2C**, **Supplementary Table S5**). Approximately 100 transcripts fit these thresholds. Several of these transcripts were highly duplicated in Eutardigrada and were also conserved in Heterotardigrada. By querying these AlphaFold2 modeled protein structures against the AlphaFold2 SwissProt database using FastSeek, several of the proteins were annotated. We found two proteins that we identified as related genes to be induced from the transcriptome analysis (**Figure 3AB**); 5-hydroxytryptamine receptor 1E (BV898_15649.t01, AF-Q6VB83-F1-model_v4.pdb E-value = 6.22E-12, RMSD between 163 pruned atom pairs is 1.124 angstroms across all 353 pairs: 10.861) and Ecdysone-induced protein 78C (BV898_05254.t01, AF-P45447-F1-model_v4 E-value 4.48E-08, RMSD between 84 pruned atom pairs is 1.044 angstroms; across all 391 pairs: 32.283). In addition, we found 18 transcripts with high IUPRED3 scores above 0.9 (four with 1.0), indicating highly disordered proteins (**Figure 3CDEF**). Only one transcript (BV898_07132.t01) showed homology to a Swiss-Prot gene (28 kDa heat- and acid-stable phosphoprotein). The Dsup gene (BV898_01301) also fit into this category, thus suggesting that these four may be novel nuclear protection / repair proteins. Additionally, we screened the 102 genes for those with known DNA binding motifs and found only two: BV898_13251 and BV898_10351. The gene BV898_13251 was found to harbor a “Helix-loop-helix DNA-binding domain”, found in TFEB transcription factors, and BV898_10351 was expressed at low levels (< TPM 10). We concluded that we could not identify potential DNA-binding protection related genes through our current dataset.

### 2.3. Metabolome analysis

To validate whether tryptophan metabolites are accumulated during anhydrobiosis entry and bleomycin response, mirroring the DEG analysis, we subjected active, anhydrobiotic, and bleomycin exposed *H. exemplaris* specimens to metabolomic analysis using liquid chromatography-mass spectrometry (LC-MS) and capillary ion chromatography-mass spectrometry (capillary IC-MS). We detected a total of 237 compounds, with relatively high Spearman correlations between replicates. PCA analysis of the metabolite profiles showed high consistency between control (active) and bleomycin-exposed (Bleomycin) samples, but with some variance in anhydrobiosis samples (Anhydrobiosis) (**Supplementary Figure 3A**). From these metabolite profiles, we detected a total of 113 compounds with significant differences between conditions (pairwise T-test, FDR < 0.05); 92, 49, and 95 compounds for Control vs Anhydrobiosis (**Figure 3**, **Supplementary Figure S3B**), Control vs Bleomycin, and Anhydrobiosis vs Bleomycin, respectively. Among the compounds that showed higher concentration in Anhydrobiosis, or Bleomycin compared to Control, eight were common to both conditions, 17 and 32 were Bleomycin and Anhydrobiosis specific. These eight compounds were composed of 3-methylhistidine, 3-phenyllactate, ADP-ribose, CDP-choline, deamido-NAD+, lysine, riboflavin, and S7P. KEGG pathway enrichment analysis of each condition combination (e.g., Control vs Anhydrobiosis, Control vs Bleomycin, Anhydrobiosis vs Bleomycin), indicated enrichment of several KEGG pathways, of which only Control vs Bleomycin showed enrichment in tryptophan metabolism (FDR = 0.002264990). Mapping of transcriptome and metabolome data to the KEGG tryptophan metabolism map (ko00380) revealed accumulation of tryptophan, 5-hydroxy-L-tryptophan, L-kynurenine and 3-hydroxy-anthranilate compounds and the regulation of enzymes contributing to this pathway (**Figure 4**, **Supplementary Figure S3C**). Interestingly, we did not see accumulation of these compounds in anhydrobiosis samples, although the corresponding enzymes seem to be regulated.

**Fig. 3.**
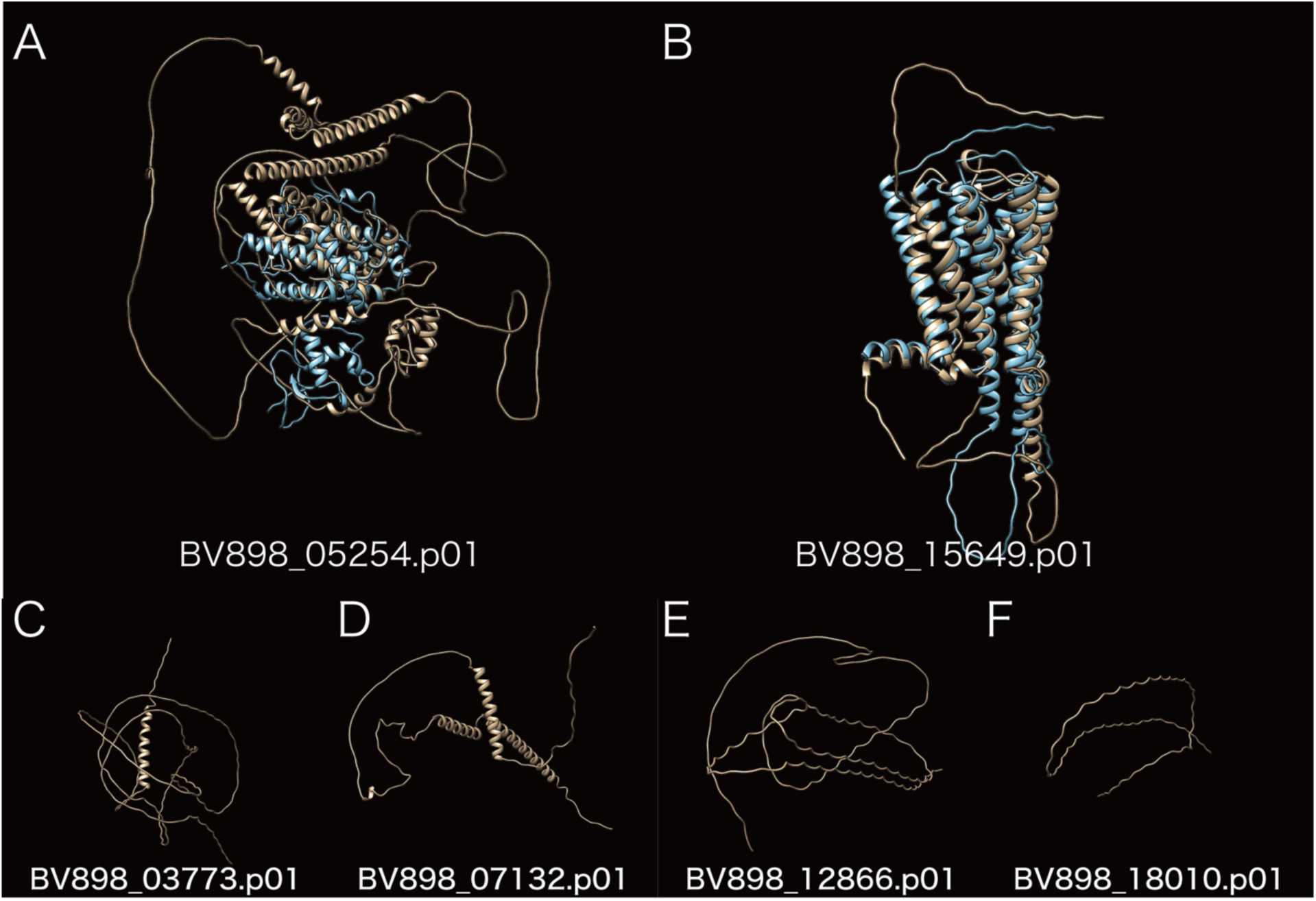
AlphaFold structures of NLS-containing proteins. AlphaFold2 structures modeled from NLS-predicted proteins that had no SwissProt BLASTP hits (E-value < 1e-15). [AB] Structures for those that had a FoldSeek hit against the AlphaFold2 SwissProt database (E-value < 1E-05). The yellow and blue structures indicate the reference and the predicted *H. exemplaris* protein structures, respectively. [A] BV898_05254.p01 (5-hydroxytryptamine receptor 1E (AF-Q6VB83-F1-model_v4.pdb E-value = 6.22E-12, RMSD between 163 pruned atom pairs is 1.124 angstroms across all 353 pairs: 10.861) [B] Ecdysone-induced protein 78C (AF-P45447-F1-model_v4 E-value 4.48E-08, RMSD between 84 pruned atom pairs is 1.044 angstroms; across all 391 pairs: 32.283) [C-F] Transcripts that had an IUPRED3 score of 1.0 [C] BV898_03773.p01, [D] BV898_07132.p01 [E] BV898_12866.p01 [F] BV898_18010.p01.

**Fig. 4.**
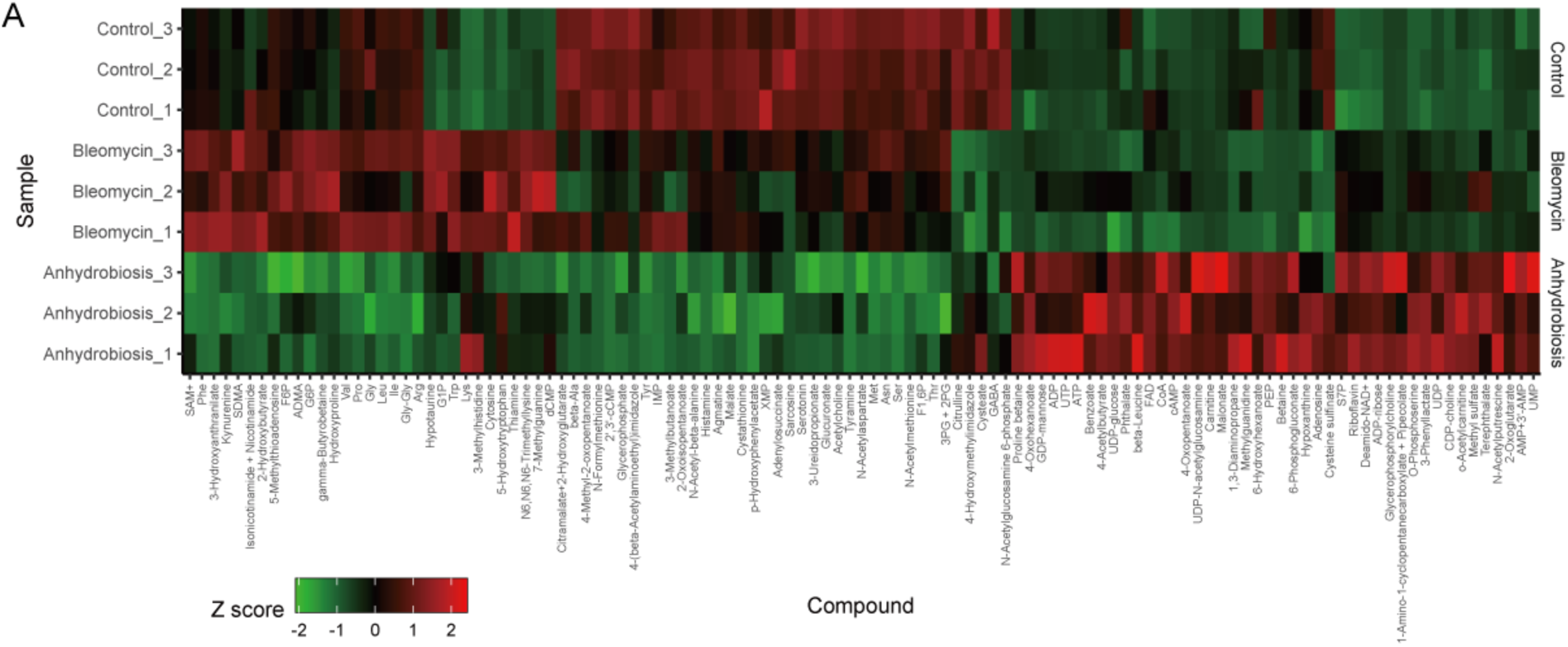
Metabolomic profiles of active, anhydrobiotic, and bleomycin-exposed *H. exemplaris*. Z-scaled concentration profiles for 113 metabolites with significant differences between Control-Anhydrobiosis, Control-Bleomycin, Anhydrobiosis-Bleomycin.

**Fig. 5.**
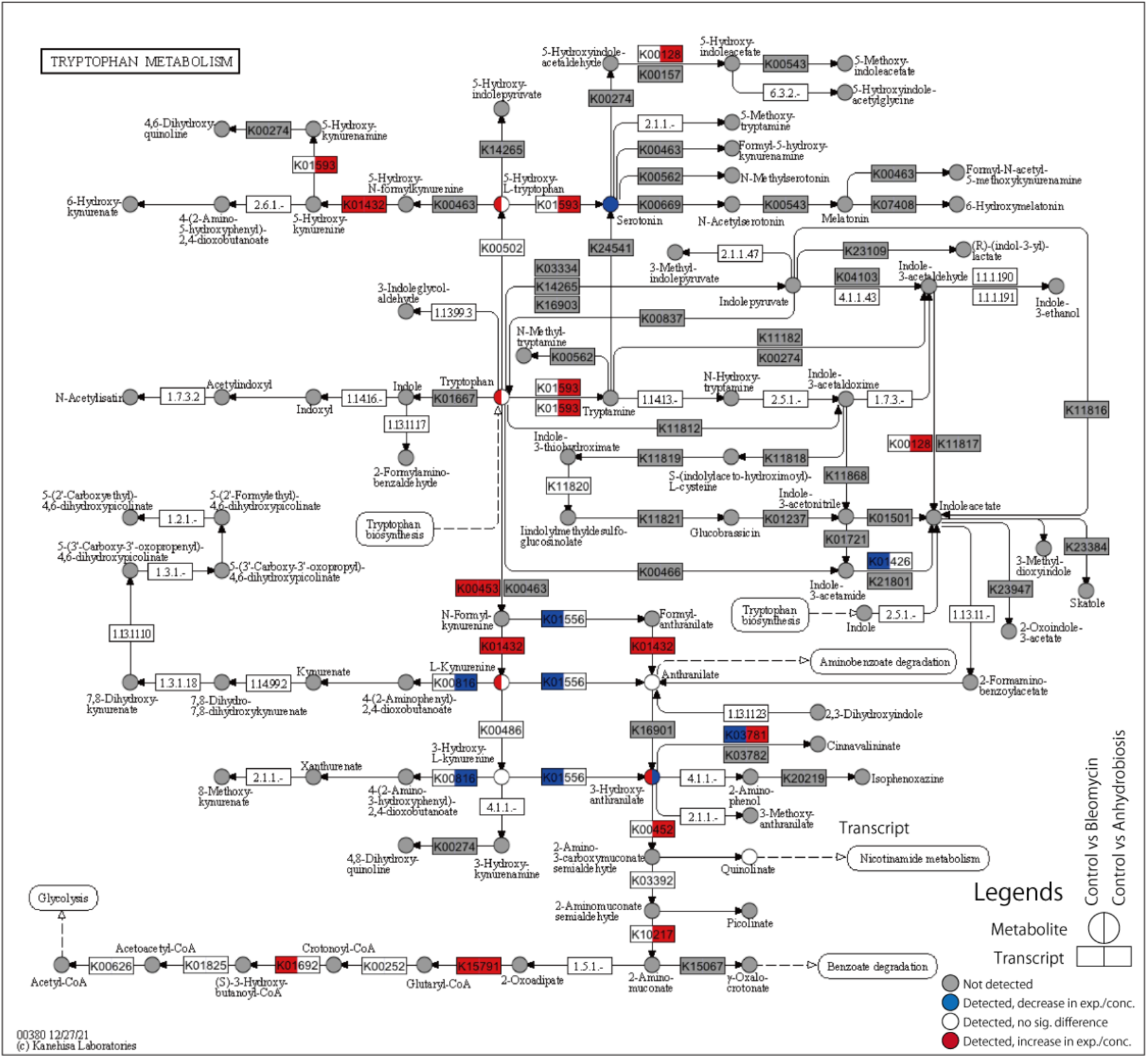
KEGG pathway map of tryptophan metabolism. Data for two comparisons are indicated in a single circle (metabolite) or square (transcript); For metabolites, Left: Control vs Bleomycin; Right : Control vs Anhydrobiosis; For transcripts; Left: Control vs Bleomycin-00h (24h exposure, no recovery time), Right: Active vs Tun. Colors correspond to the following conditions: Gray: not detectable or not detected (0µg/µL for each condition), Blue: decrease in expression/concentration (lower in Bleomycin/Anhydrobiosis); White: detected, but no significant difference; Red: increase in expression/concentration (higher in Bleomycin/Anhydrobiosis).

## 3. Discussion

Tardigrades in general have been observed to have high radiation tolerance, which has been hypothesized to be a byproduct of the acquisition of the anhydrobiosis mechanism. We have previously performed transcriptome analysis of *R. varieornatus* exposed to gamma rays and UVC to identify core components contributing to this cross-tolerance ^33)^. Building on this, we tested for such factors that could be induced by a more chemical approach, in this study using bleomycin. Bleomycin causes both direct and indirect DNA damage,, making it comparable to previous radiation-based methods that mainly cause indirect damage through reactive oxygen species.

In this study, we used the bleomycin concentration of 100µM. In Chinese hamster DC-3F fibroblasts, this concentration is approximately equivalent to 1,500,000 Gy (1nM to 15Gy) ^40)^. However, considering that (1) bleomycin itself does not diffuse through the plasma membranes, (2) we used individual tardigrades, which is quite different from exposure to cell cultures, (3) tardigrades have cuticular exoskeleton that may further impede cellular uptake, 100µM would not directly correspond to this extremely high gamma ray dose. A concentration of 50 µM has been used in *Caenorhabditis elegans* ^41)^, 10-50 µg/mL in *Drosophila melanogaster* (corresponding to 7-35 µM for 1415.55g/mol bleomycin) ^42, 43)^, so considering the higher tolerance to DNA damage observed in *H. exemplaris*, the concentration used in our study is neither extremely high nor low. A previous study used an exposure of 100µM for four or five days against *H. exemplaris* and analyzed the transcriptomic response at the point of exposure end. Therefore, in order to induce DNA damage, but achieve near 100% survival, *H. exemplaris* specimens were exposed to 100 µM bleomycin for 24h and were sampled over the time course of 24 h after removal of tardigrades from bleomycin solution.

We generated time-course transcriptome sequencing data on *H. exemplaris* specimens recovering from a 24h exposure to 100µM bleomycin. We observed reduction in movement during the bleomycin exposure, similar to that observed in UVC-exposed *R. varieornatus* ^33, 44)^. We observed a dynamic shift in the transcriptome at 0-1h and 9-24h after bleomycin clearance, suggesting the existence of an early, middle, and late response. While this three-step response was similar to that observed in *R. varieornatus* upon irradiation, the timing of the transcriptome shifts was relatively different; 4-6h and 24-36h for UVC, 6-9h in gamma-ray response ^32, 33)^. This may be due to several factors; the early response may be induced during the 24h bleomycin exposure, differences in the requirement of stress response factors between *R. varieornatus* and *H. exemplaris*, differences in the mechanism by which gamma-ray, UVC, and bleomycin cause DNA damage.

Clustering of the transcriptome profiles and detection of enriched gene ontologies revealed that mismatch repair (MMR) and nucleotide excision repair (NER) are induced in the early response (Groups 3 and 8) and non-homologous end joining (NHEJ) in the middle response (Group 5). This suggests that *H. exemplaris* may repair minor damage during drug exposure, followed by the repair of devastating double strand breaks after drug clearance. The fact that cell replication is rare in active *H. exemplaris* ^45)^, with the exception of the digestive tract, and that they are equipped with specialized protection mechanisms such as Dsup, suggests that double strand breaks may not be a priority in damage repair, but rather a focus on minor DNA damage that would cause disruption of gene expression. Further analysis in the early, middle, and late responses showed that DNA replication and central carbon metabolism are regulated in the late response, thus suggesting that DNA repair and cell replication may occur at least 12-24 hours after exposure to bleomycin. In addition, we found regulation of mitochondria-related genes during the middle response, suggesting mitochondrial reorganization at this stage. An ultrastructural study in *H. exemplaris* anhydrobiosis has observed abnormal morphology in the mitochondria ^46)^, additionally regulation of the mitochondrial chaperone BCS1 has also been observed ^31)^, both suggesting the presence of mitochondrial stress.

We also validated the expression of previously identified anhydrobiosis genes and observed the induction of CAHS, AMNP, and Rv.g241 orthologs. These gene families were also found to be shared between anhydrobiosis, bleomycin recovery, and UVC response in *R. varieornatus*. In addition, a single AMNP ortholog (BV898_01090) and Rv.g241 ortholog (BV898_18007) were also found to be significantly induced in the proteomic data of D942 exposed *H. exemplaris* ^47)^, suggesting regulation of this ortholog through AMPK ^48)^ and highlighting the importance of these two gene families in the stress response in *H. exemplaris* and *R. varieornatus*. AMNP and Rv.g241 gene families were identified as stress responding families through transcriptome analysis in UVC exposed *R. varieornatus* ^33)^. AMNP gene family was identified as a Mn^2+^ dependent peroxidase, while the function of Rv.g241 remains to be identified, but considering that it is conserved at the C-terminus of bacterial peroxidases, it may be some kind of cofactor that enhances the antioxidative functions of peroxidases. Surprisingly, we found that many CAHS orthologs are also induced. CAHS proteins have been found to form a high-order structure in a desiccated state, which has been hypothesized to protect cellular molecules ^49)^. Such high-order structures would not be required during recovery from damage; therefore, CAHS proteins may have additional stress-responsive functionalities, presumably related to its disordered nature with highly abundant charged residues. A recent study employing transcriptome sequencing on bleomycin exposed *H. exemplaris* has identified a DNA damage response gene TDR1 (BV898_14257) ^39)^. While the TDR1 gene was highly induced in our bleomycin dataset similar to the previous study, we did not observe induction in the anhydrobiosis course. This may imply that there is limited DNA damage during anhydrobiosis, or the induction of TDR1 requires prolonged recovery period. We hypothesized that the *R. varieornatus* TDR1 ortholog may be induced in our previous gamma ray exposure transcriptome dataset ^32)^; however, we did not find a TDR1 ortholog in other accessible tardigrade genomes using BLASTp using the truncated BV898_14257 amino acid sequence (E-value < 1E-5)^33)^. Synteny based analysis suggested g12065.t1 as a candidate ortholog with no known functional domain annotation, however, the sequence similarity was extremely low, and AlphaFold2 prediction suggested opposite alpha-/beta- propensities. The expression of Rv.g12065 was moderate (approx. TPM 50) for the gamma ray exposure time course. These data may imply that this gene may be a *Hypsibius* specific gene, or an ortholog pair with extremely low similarity, as observed between HeDsup and RvDsup. We have also identified approximately 100 novel genes that encode contain a nuclear localization signal suggestive of a nucleolus-localizing protein. Many of these proteins are highly disordered, a feature similar to tardigrade-specific anhydrobiosis genes ^22, 37, 38)^. These proteins would be future candidates for future functional analysis.

We then compared the transcriptome response to anhydrobiosis entry. We found that transcripts regulated during anhydrobiosis entry shared the most transcripts with the early response (42%), followed by the late (25%) and middle (16%) responses. Gene Ontology enrichment analysis identified 37 gene ontology terms to be enriched in the early response-anhydrobiosis gene set. This list included many terms related to tryptophan metabolism and the oxidation-reduction process. Tryptophan metabolism has recently emerged as an important pathway in human therapeutics, immunology, and neuronal function ^36)^. A metabolome study in *Polypedilum vanderplanki*, an anhydrobiotic insect, has identified the tryptophan oxidation pathway as one of the most anhydrobiosis-responsive pathways, *e.g.*, kynurenic acid and xanthurenic acids showed high accumulation during anhydrobiosis recovery ^50)^. In our metabolomic analysis, we observed accumulation of tryptophan metabolites in only in bleomycin-exposed samples, not during anhydrobiosis entry. This is not consistent with the gene expression profiles; most enzymes in the tryptophan metabolism pathway are induced in both bleomycin exposure and anhydrobiosis entry. Comparison of the tryptophan metabolite profiles with *P. vanderplanki* indicated similar profiles; constant 3-hydroxy-kynurenine concentration, no significant difference in expression of kynurenine aminotransferase (K00816) during bleomycin exposure (down-regulated during anhydrobiosis entry). We did not detect xanthurenic acid and kynurenic acid in all samples. These data suggest that while tryptophan metabolism may be an important factor in tardigrades and *P. vanderplanki*, there are differences in how these pathways contribute to anhydrobiosis itself.

In this study, we have performed time-course transcriptome analysis of bleomycin-exposed *H. exemplaris* to identify factors that constitute the core element of the cross-tolerance machinery. Through comparative transcriptomics, we propose that previously identified stress responders as well as the tryptophan metabolism are such elements. We attempted to validate this through metabolomic analysis, suggesting different profiles between anhydrobiosis and chemical stress. Taken together, we propose that antioxidative functions through tryptophan metabolism may play a role in the tardigrade stress response, to what extent remains to be clarified.

## 4. Methods

### 4.1. Tardigrade culture

*H. exemplaris* Z151 strain was cultured following our previous studies ^31, 34)^. Briefly, specimens were placed on 2% agar gels using Volvic water, supplied with Volvic water containing *Chlorella vulgaris*. Individuals were collected and moved to a new agar plate every 7 days.

### 4.2. Transcriptome sequencing of bleomycin exposed individuals

Fifty individuals per sample were placed in 96 well flat bottom plates and was exposed to 200µL 100µM bleomycin (Funakoshi, Cat No. B4518, dissolved in Volvic water) for 24 hours (N=5). A control series exposed to Volvic water was also set. After 24 hours, the control specimens and T0 specimens were collected and placed in a microtube with the least amount of water possible and frozen at -80°C. The remaining samples were washed twice with 300µL Volvic water to remove bleomycin and were placed on 45mm agar plates with 2mL Volvic water supplemented with 10µL *Chlorella* for feeding. These specimens were incubated at 18°C until sampling. After 1,3, 6, 9, 12, and 24 hours, each sample was collected as described above and frozen at -80°C.

Frozen samples were submitted for transcriptome sequencing (N=3). Total RNA was extracted using Direct-Zol Micro Kit and quantified with Qubit RNA HS. An Illumina sequencing library was constructed from 30ng of total RNA using NEB Next Ultra RNA according to the manufacture’s protocol. The final library was sequenced on the NextSeq 500 instrument (Illumina). Multiplexed reads were demultiplexed with bcl2fastq (Illumina). The quality of RNA-Seq reads was validated using FastQC v0.11.3 ^51)^.

### 4.3. Informatics analysis

The genomes, coding and amino acid sequences and annotations for *H. exemplaris* nHd3.1.5 and *R. varieornatus* v101 were downloaded from http://ensembl.tardigrades.org ^31)^. In addition, we predicted nuclear localizing proteins using NLStradumus v1.8 ^52)^. Sequenced reads were mapped to CDS and quantified with RSEM v1.2.26 ^53)^ and bowtie2 v2.2.9 ^54)^ using the Trinity align_and_estimate_abundance.pl utility ^55)^. Differential expression was tested with DESeq2 v1.22.2 ^56)^ using the Trinity run_DE_analysis.pl, and transcripts with adjusted *p*-values below 0.05 and fold change over 1.5 or below 2/3 were designated as differentially expressed genes. The transcriptome profile was clustered by the Ward method using Spearman correlation. Each DEG cluster was subjected to Gene Ontology enrichment analysis with GOstat v2.48.0 ^57)^ using InterProScan Gene ontology annotations ^58)^.

To determine gene families that are commonly regulated in multiple stresses, we used the transcriptome sequencing data of *H. exemplaris* anhydrobiosis (GSE94295) ^31)^ as well as the time-course RNA-Seq data of UVC-exposed *R. varieornatus* (GSE152753) ^31)^. The transcriptome sequencing data of *H. exemplaris* were subjected to differential expression analysis using the same method described above. DEGs of *R. varieornatus* UVC exposure were obtained from our previous study^33)^. To identify ortholog families, the amino acid sequences of *H. exemplaris*, *R. varieornatus*, and other tardigrade genomes from our previous analysis ^33)^ were clustered using OrthoFinder v2.5.1 ^59)^. Ortholog clusters containing DEGs of *H. exemplaris* bleomycin exposure, anhydrobiosis entry, and *R. varieornatus* UVC exposure were detected, and the annotations were validated. For gene families that were not previously annotated, we submitted the amino acid sequences to ColabFold (Alphafold_batch https://colab.research.google.com/github/sokrypton/ColabFold/blob/main/batch/Alp <u>haFold2_batch.ipynb</u>) ^60)^ and predicted the protein structure with AlphaFold ^61)^. Only one model was predicted for amino acid sequences capable of prediction. The predicted protein structures were submitted to DALI v5 search ^62)^ or FoldSeek v6.29e2557 ^63)^ against the Alphafold Protein Structure Database ^64)^. Homology were determined by (1) identity > 30 or (2) z-scale > 10 for DALI and E-value < 1e-5 for FoldSeek searches. Tardigrade predicted proteomes were screened for TDR1 orthologs using BLASTp search (E-value < 1e-5). Additionally, we conducted synteny searches using JCVI McScan between *R. varieornatus* and *H. exemplaris*^65)^.

### 4.4. Metabolome analysis

Hydrophilic metabolites were analyzed using two different analytical methods. Cationic metabolites were measured using liquid chromatography-mass spectrometry (LC-MS), as described previously ^66, 67)^. Briefly, LC-MS analysis was performed using an Agilent 1290 Infinity LC (Agilent Technologies, Santa Clara, CA) equipped with a Q Exactive Orbitrap mass spectrometer (Thermo Fisher Scientific, San Jose, CA). Separations were performed on a HILIC-Z column (150 × 2.1 mm, 2.7 µm; Agilent Technologies). The mobile phase was composed of 20 mM ammonium formate + 0.25% (*v/v*) formate (A) and 20 mM ammonium formate + 0.25% (*v/v*) formate in 90% (*v/v*) acetonitrile (B). The flow rate was 0.25 mL/min, and the following linear gradient was used: 0–15 min, 100% to 70% B; 15–20 min, 70% to 10% B; 20–23 min, 10% B, followed by equilibration with 100% B for 7 min. The injection volume was 1 µL, and the column temperature was maintained at 40 °C. The Q Exactive mass spectrometer was operated in a heated electrospray ionization (HESI) positive-ion mode using the following source parameters: spray voltage = 3.5 kV, capillary temperature = 250 °C, sheath gas flow rate = 40 (arbitrary units), auxiliary gas flow rate = 10 (arbitrary units), auxiliary gas temperature = 300 °C, sweep gas flow rate = 0, S-lens = 35 (arbitrary units). Data were acquired using a combination of full MS scan mode and parallel reaction monitoring (PRM) mode. The parameters in full MS scan mode were as follows: resolution, 35,000; auto gain control target, 3 × 10^6^; maximum ion injection time, 200 ms; scan range, 50–700 *m/z*. The parameters in PRM mode were as follows: resolution, 17,500; auto gain control target, 2 × 10^5^; maximum ion injection time, 100 ms; inclusion *m/z* list: 104.0706 (GABA), 118.0863 (Val), 132.1019 (Ile, Leu), 166.0863 (Phe), 182.0482 (methionine sulfone).

Anionic metabolites were analyzed by capillary ion chromatography-mass spectrometry (capillary IC-MS) as previously described^68)^. Briefly, capillary IC-MS was performed using a Dionex ICS-5000+ system coupled with a Q Exactive Orbitrap mass spectrometer. Separation of anionic metabolites were performed using a Dionex IonPac AS11-HC-4 µm column (250 × 0.4 mm, 4 µm; Thermo Fisher Scientific). The injection volume was 0.4 µL, and the column temperature was maintained at 35 °C. The eluent flow rate was 20 µL/min and the potassium hydroxide (KOH) gradient was as follows: 0-2 min, 1 mM; 2-16 min, 1 mM to 20 mM; 16-35 min, 20 mM to 100 mM; 35-40 min, 100 mM; followed by equilibration with 1 mM KOH for 5 min. Isopropanol containing 0.1% acetate was delivered as the make-up solution at 5 µL/min. The Q Exactive mass spectrometer was operated in ESI negative-ion mode. ESI parameters were as follows: sheath gas, 20 (arbitrary units); auxiliary gas, 10 (arbitrary units); sweep gas, 0; spray voltage, 4.0 kV; capillary temperature, 300 °C; auxiliary gas temperature = 300 °C; S-lens, 50 (arbitrary units). Data were acquired in full MS scan mode and the parameters were as follows: resolution, 70,000; auto gain control target, 3 × 10^6^; maximum ion injection time, 100 ms; scan range, 70–1,000 *m/z*.

Quantified metabolites were further normalized by the total amount of compounds detected to the average of all samples. Metabolite profiles were subjected to pairwise T-test in R, and compounds with FDR < 0.05 were considered significant differences. Compound names were converted to KEGG IDs using MetaboAnalyst 5.0 ^69, 70)^, and those that did not have a correct match were further searched in PubChem, ChEBI, or KEGG databases. The corresponding compounds could not be found in any of the databases, or were a peak containing two compounds that could not be separated with our setup, and were therefore omitted from enrichment analysis: “1-Amino-1-cyclopentanecarboxylate + Pipecolate”, “3PG + 2PG”, “4-Acetylbutyrate”, “4-Hydroxymethylimidazole”, “4-Oxohexanoate”, “4-Oxopentanoate”, “AMP+3’-AMP”, “Citramalate+2-Hydroxyglutarate”, “Isonicotinamide + Nicotinamide”, “SDMA”. Linking data between KEGG compound IDs and pathway IDs were obtained from the KEGG REST service for all analyzed metabolites. For each KEGG pathway, we preformed Fisher’s exact test to detect enrichment using FDR-corrected p-values < 0.05 as the threshold. Metabolite and transcriptome profiles were visualized on KEGG pathway maps with Pathview^71)^.

### 4.5. Data analysis

Statistics and data manipulation were performed with R and custom scripts using the G-language Genome Analysis Environment v.1.9.1^72, 73)^. Biovenn was used to visualize area -proportional Venn diagrams ^74)^.

### 4.6. Data availability

RNA-Seq reads have been uploaded to GEO under the accession ID SE168917.

## Supplementary Table legends

Supplementary Table S1. Summary of RNA-Seq data produced in this study. Supplementary Table S2. Gene ontology enrichment analysis for each of the eight groups.

Supplementary Table S3. Gene ontology enrichment analysis for genes common between anhydrobiosis and each response group (Early, Middle, Late).

Supplementary Table S4. Annotation of stress-responsive gene families induced in at least two conditions in anhydrobiosis, bleomycin, and Rv-UVC. Yellow and green rows indicate tardigrade specific gene families and those conserved in at least one non-tardigrade or Metazoa analyzed. Conservation profiles, number of orthologs induced in stress conditions, protein annotation from nHd.3.1.5, DALI hits from Alphafold structure database (I>30 and Z>10).

Supplementary Table S5. NLS containing proteins that are induced in anhydrobiosis and either one of the response groups (Clusters A/B/C). NLS was predicted with NLStradamus.

## Supplementary Figure legend

**Supplementary Fig. S1.**
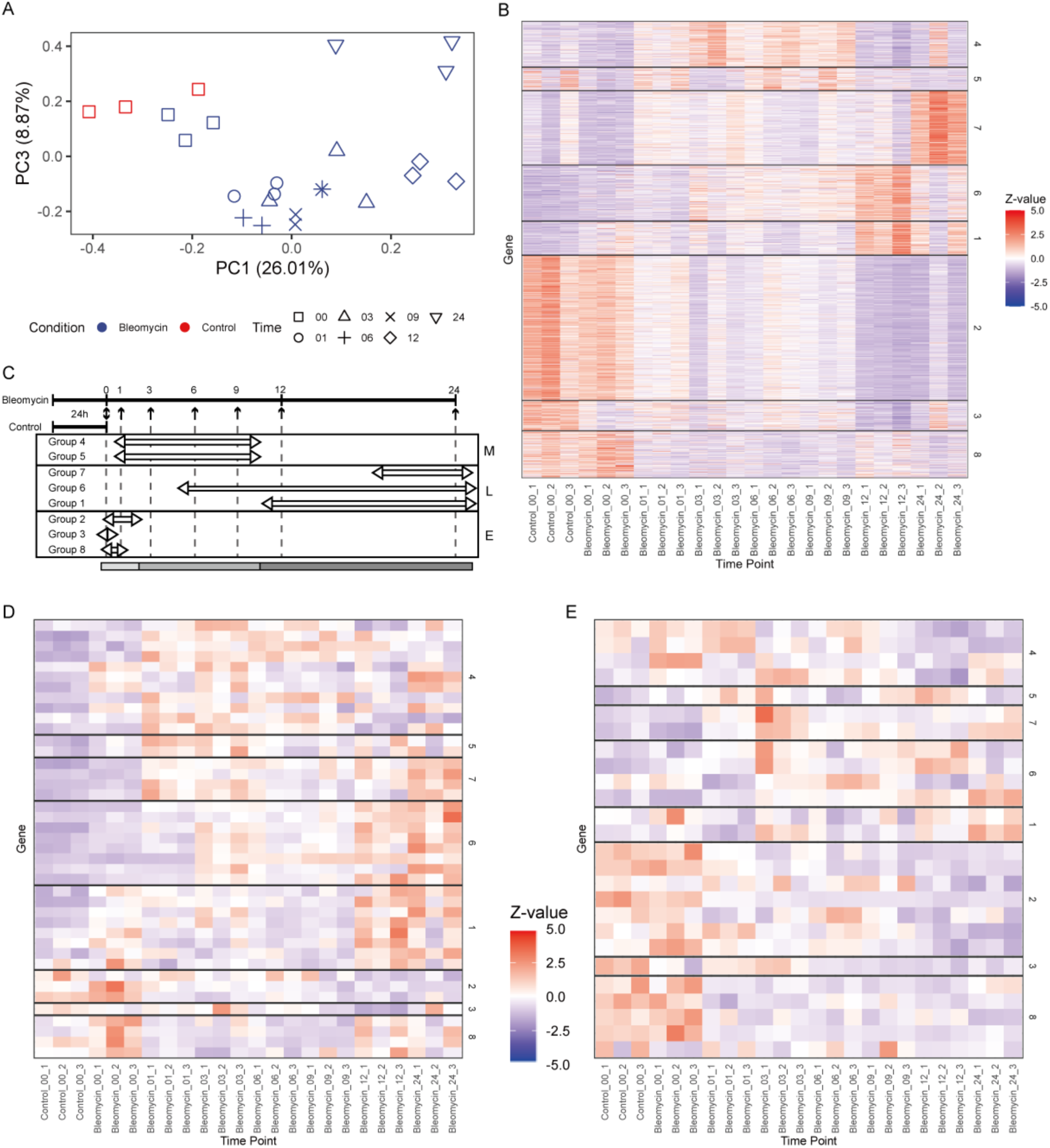
[A] PCA of transcriptome profiles. Transcripts with average expression > 1 were subjected to PCA. [B] Clustering of expression profiles. Transcripts with average expression > 1 were clustered based on Spearman’s correlation and split into eight groups based on total within sum of squares. A dynamic change was observed between 0-1h, 9-12h, 12-24h. [C] An illustration of the broader clustering of the eight groups. E, M, and L indicate the Early, Middle, and Late responses. [DE] Expression profiles of previously identified novel tardigrade-specific stress responsive-proteins. [D] AMNP family [E] Rv.g241 gene family.

**Supplementary Fig. S2.**
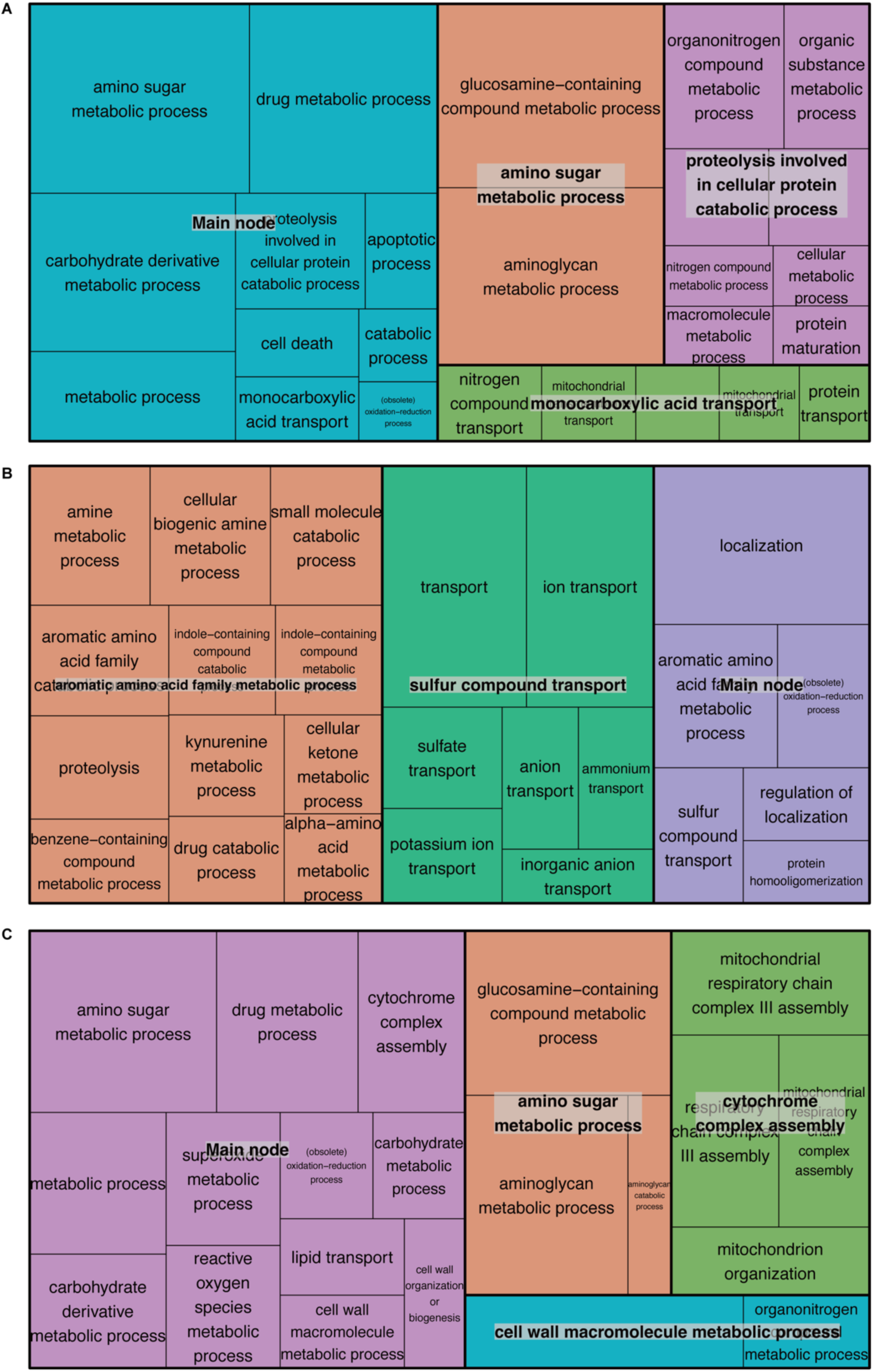
Treemap of Revigo-analysis on Gene Ontology enrichment analysis of each group. [A] Cluster Early [B] Cluster Middle [C] Cluster Late.

**Supplementary Fig. S3.**
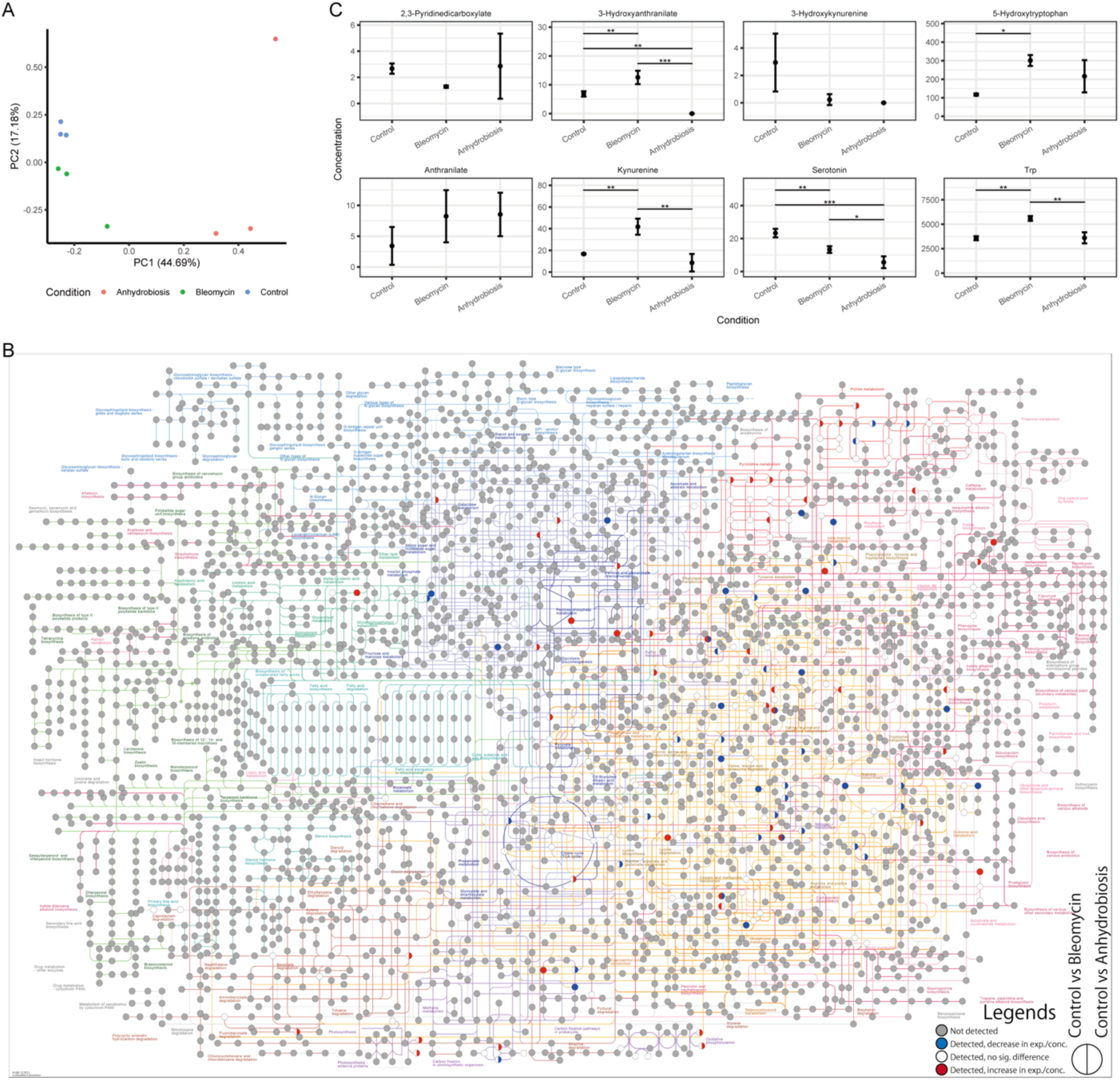
General analysis on metabolome data. [A] PCA of metabolome compound concentrations. [B] KEGG pathway colored with metabolite profiles. Data for two comparisons are shown in a single circle Left: Control vs Bleomycin; Right : Control vs Anhydrobiosis. Colors correspond to the following conditions: Colors correspond to the following conditions: Gray: not detectable or not detected (0µg/µL for each condition), Blue: decrease in expression/concentration (lower in Bleomycin/Anhydrobiosis); White: detected, but no significant difference; Red: increase in expression/concentration (higher in Bleomycin/Anhydrobiosis) [C] Concentration profiles for trytophan metabolites.

## Acknowledgements

We thank Naoko Ishii, Ayako Shirahata, and Yuki Takai for the tardigrade experiments and transcriptome sequencing. We also thank Satsuki Ikeda for the metabolome analysis. *C. vulgaris* used to feed the tardigrades was provided courtesy of Chlorella Industry. This work is supported by KAKENHI Grant-in-Aid for Transformative Research Areas (A) from the Japan Society for the Promotion of Science (JSPS, grant Number 21H05279), Joint Research by Exploratory Research Center on Life and Living Systems (ExCELLS program Nos. 19-501 and 22EXC601) and partly by research funds from the Yamagata Prefectural Government and Tsuruoka City, Japan.

